# Mechanisms Underlying Allosteric Modulation of Antiseizure Medication Binding to Synaptic Vesicle Protein 2A (SV2A)

**DOI:** 10.1101/2025.05.05.652227

**Authors:** Anshumali Mittal, Matthew F. Martin, Laurent Provins, Adrian Hall, Marie Ledecq, Christian Wolff, Michel Gillard, Peter S. Horanyi, Jonathan A. Coleman

## Abstract

Brivaracetam (BRV) and levetiracetam (LEV) are antiseizure medications (ASMs) that target synaptic vesicle protein 2A (SV2A), while UCB1244283 acts as a positive allosteric modulator of these medications. The SV2A-BRV-UCB1244283 complex reveals how UCB1244283 allosterically enhances BRV binding by occupying an allosteric site near the primary binding site, preventing BRV dissociation. This allosteric site, formed by hydrophobic and uncharged residues, is a novel small-molecule binding site in SV2A. Structural analysis and mutagenesis suggest that an allosteric network between the primary and allosteric sites governs high-affinity ASM binding. UCB1244283 selectively binds SV2A over SV2B and SV2C, with specific mutations disrupting binding. The structure explains why UCB1244283 binding to SV2A selectively allows interaction with specific ASMs but not others due to steric hinderance. Structural comparison reveals that distinct conformational differences between the SV2A-BRV-UCB1244283 and other SV2A-ligand complexes, particularly in the transmembrane domain, influence binding at both sites. Future research will explore potential therapeutics targeting the allosteric site and their impact on SV2A regulation.

**Teaser:** UCB1244283 enhances LEV and BRV binding to SV2A, revealing an allosteric site that could aid in developing targeted therapeutics.

## Introduction

Epilepsy is a common neurological disorder characterized by an abnormal activity of neurons which leads to seizures. Approximately 1% of the population will develop a seizure disorder, which can cause exceptional challenges for affected individuals^1^. Individuals with unresponsive seizures face a lifetime of physical, mental, and social challenges as well as mortality rates more than three times higher than the rest of the population^2^. A large number of antiseizure medications (ASMs) are approved as therapeutics for different seizure types, however, 30% of patients cannot achieve seizure control with current available medications either due to poor efficacy or undesirable side-effects^3^. Therefore, there is a pressing need to better understand the underlying etiologies of epilepsies and to develop more targeted treatments. Levetiracetam (LEV, Fig. 1a) is distinct from other ASMs due to its unique mechanism of action, involving specific binding to synaptic vesicle protein 2A (SV2A)^4^. Since its FDA approval in 1999, LEV (tradename: Keppra) has emerged as a highly effective ASM for the treatment of treatment of myoclonic and generalized tonic-clonic seizures^5,6^. LEV and related compound brivaracetam (BRV, Fig 1a), have been shown to reduce glutamatergic transmission, a mechanism that is critical for the generation of seizures^7,8^. SV2A belongs to the major facilitator superfamily (MFS) of neuronal transporters and includes SV2B and SV2C^9–13^. LEV and BRV bind selectively to SV2A with at least 100-fold higher affinity compared to SV2B and SV2C^3,5,14,15^. Padsevonil (PSL) was rationally designed to bind all SV2 members (SV2s) with nanomolar affinity and was previously investigated in clinical trials^15,16^. The luminal domain (LD) of SV2s serves as a receptor for mediating neuronal entry of exogenous toxins, such as botulinum toxin (BoNT) and tetanus toxin, through endocytosis^17–19^. BoNT is used clinically to treat conditions like eyelid twitching, neck muscle spasm and cosmetically for facial lines^20^.

**Fig. 1.**
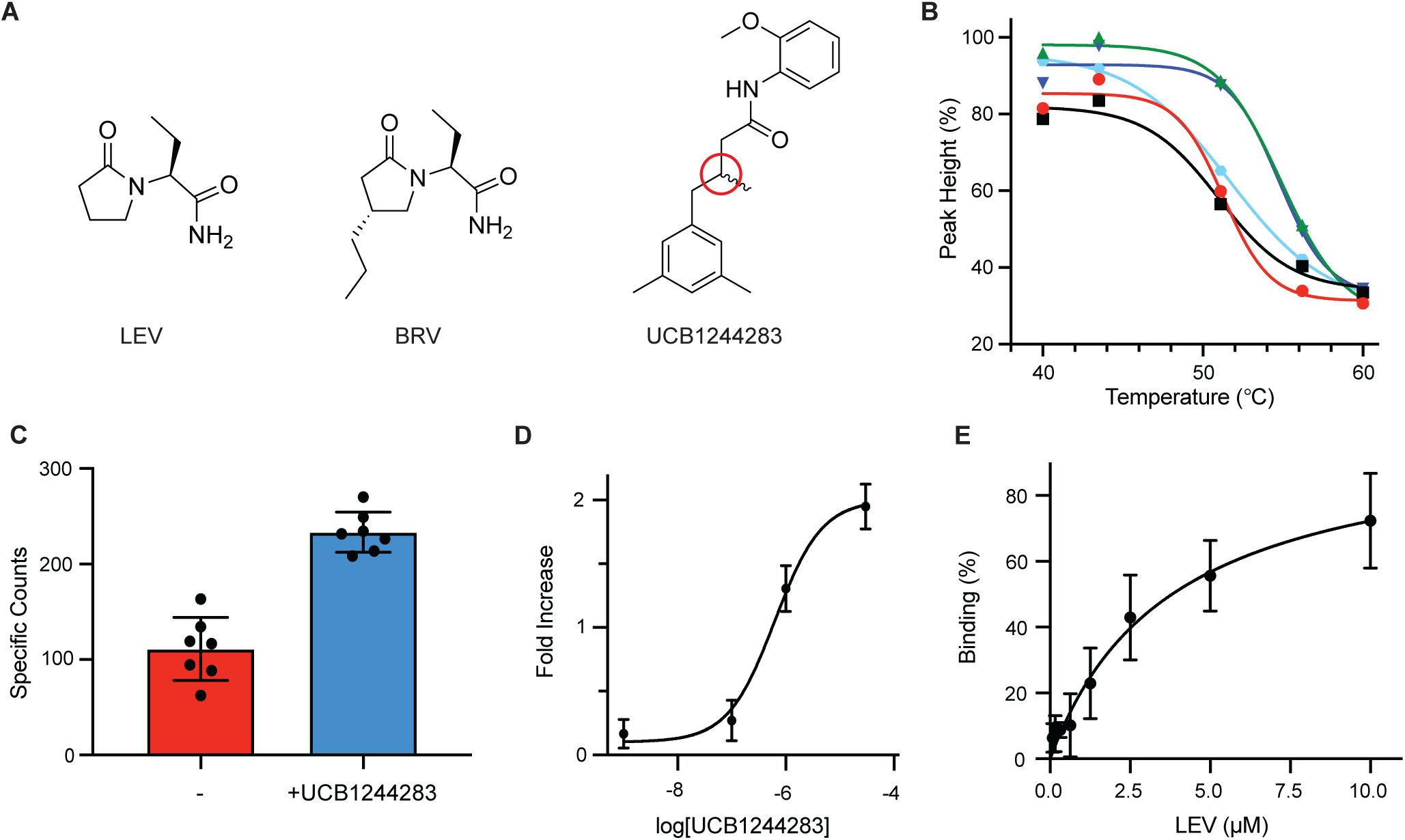
UCB1244283 exerts an allosteric effect on the SV2A-LEV complex. **A.** Chemical structures of SV2A-specific ligands, Levetiracetam (2S-(2-oxo-1-pyrrolidinyl)butanamide), Brivaracetam ((2S)-2-[(4R)-2-oxo-4-propylpyrrolidin-1-yl]butanamide), and UCB1244283 (4-(3,5-dimethylphenyl)-N-(2-methoxyphenyl)-3-methylbutanamide), chiral center is shown with red circle. **B.** Thermostability of mVenus-tagged SV2AΔ64 with no ligand (red), 20 µM BRV (green), 30 µM UCB1244283 (black), BRV-UCB1244283 (blue) and LEV-UCB1244283 (cyan). 100 µM LEV was used for the LEV-UCB1244283 experiment. **C**. UCB1243283 (30 µM) increases the maximum specific binding (B_max_) of ^3^H-LEV (∼ 4 µM) to SV2A. **D**. Concentration-response curve for measuring EC_50_ of UC1244283 (30 µM, 1 µM, 100 nM and 1 nM) by binding of ^3^H-LEV (∼ 4 µM) to SV2A. The EC_50_ was calculated 0.6 ± 0.1 µM by non-linear regression using a logarithmic concentration response sigmoidal curve. Data is shown as mean **±** SEM (n = 7) **E**. ^3^H-LEV saturation binding experiments for measuring LEV binding affinity to SV2A (*K_d_* = 3.8 ± 0.8 µM) in presence of UCB1243283. Data is shown as mean **±** SEM (n = 8).

Recently, we determined single-particle cryo-electron microscopy (cryo-EM) structures of SV2A bound to UCB-2500, a PSL-related ligand, and SV2B bound to PSL. Our reconstructions revealed detailed insights into the architecture of the transmembrane domain (TMD), LD, and intracellular domain (ICD) along with density features at the primary binding site corresponding to these ligands^21^. At the same time, four other groups have also determined structures of SV2A in complex with LEV and BRV and SV2B in an apo state^22–26^. The structures of SV2A bound with LEV or BRV closely resemble our reconstructions, except for the conformation of the lumenal half of TM1 which adopts a more closed conformation in our structure. The SV2A and SV2B TMDs exhibit signature structural motifs of MFS transporters, including a canonical MFS fold composed of twelve TM segments organized into two six-TM bundles, referred to as N-(TM1-6) and C-terminal TMDs (TM7-12). Each of these domains is formed by two structurally inverted three-TM repeats. The first helices from each three-TM repeat, TMs 1, 4, 7 and 10, are positioned at the center of the TMD to form the primary binding site, harboring the density for ASMs. SV2A and SV2B adopt a conformation resembling a lumenal-facing occluded transporter when bound to UCB-2500 and PSL^21^. In this conformation, a dityrosine motif composed of Tyr461/404 and Tyr462/405 in TM7 of SV2A/SV2B serves as the ‘lower’ lumenal gate, which is closed in our structures, preventing dissociation of the bound ASMs. However, the upper region of this cavity remains accessible from the lumenal side of SV2A and SV2B. Access to the primary ligand binding site from the intracellular side of the protein is blocked by the intracellular helix bundle and large residues in the primary binding site. In the primary binding site, two highly conserved tryptophan residues^27^, Trp300, and Trp666 in SV2A are involved in recognition of the pyrrolidone of racetams and a protonated aspartate residue, Asp670, is involved in binding to the amide of LEV and BRV and thiadiazole ring of UCB-2500 and PSL^21–23^.

Despite these advances, the structural and biochemical understanding of the SV2 ligand binding sites, including the mechanisms that regulate binding, is still limited. Previous studies identified an allosteric modulator, UCB1244283, (Fig. 1a) which binds to a different site and modulates the binding of LEV and BRV^28–30^. The administration of UCB1244283 in sound-sensitive mice has been shown to provide protection against both tonic and clonic seizures^28^. UCB1244283 does not bind to the PSL-bound SV2A complex^15^, however, incubation of UCB1244283 with SV2A results in a 2.5 or 10-fold increase in binding affinity for LEV or BRV, respectively, by slowing the rate of dissociation. Additionally, UCB1244283 increases the maximum binding capacity (B_max_) by 2-fold for LEV and 1.4-fold for BRV^29^. Interestingly, experiments using human brain membranes have shown that UCB1244283 only increases the B_max_ for LEV, while it increases both the B_max_ and binding affinity for BRV^29^. These differential effects of modulator led to the hypothesis that UCB1244283 acts differently on the BRV- and LEV-bound SV2A complexes and may expose a second LEV binding site in the SV2A-LEV complex^28^. Altogether, these studies demonstrate that there is an allosteric binding site in SV2s that modulates ligand binding to the primary site and displays distinct chemical specificity. However, the location of the allosteric site, the selectivity of UCB1244283 toward other SV2s, the possibility of a second LEV binding site, and the structural basis for why UCB1244283 does not bind to the larger ligand complexes like PSL are not known.

Here we report structural and functional analysis of UCB1244283 binding to SV2A using cryo-EM and binding experiments. Our studies provide an explanation for how allosteric ligand binding slows dissociation of primary site ligands and increases binding affinity. UCB1244283 sits directly in the luminal pathway that leads to the primary binding site and induces a conformation of TM1 that further blocks both binding sites. Our studies also uncover the underlying basis of selectivity of UCB1244283 for SV2s and other ligands like PSL. Additionally, they suggest that the distinct effects of UCB1244283 on the BRV- and LEV-bound SV2A complexes are unlikely to involve the presence of a second LEV binding site.

## Results

### UCB1244283 modulates the SV2A-LEV complex through an allosteric mechanism

We focused on functional characterization of SV2A, using LEV and BRV, which bind to the primary site of SV2A, and UCB1244283 that binds to an unidentified allosteric site (Fig. 1a). The SV2A allosteric modulator UCB1244283 is known to bind to SV2A in presence of smaller racetam ligands, such as LEV and BRV, but not to PSL^15^. These interactions appear to induce conformational changes in SV2A^15,29^. We, therefore, started by investigating the thermostability of SV2A-UCB1244283 complexes in presence of LEV or BRV to identify complexes which are suitable for subsequent structural and functional studies (Fig. 1b). The BRV-UCB1244283 bound SV2A complex exhibits enhanced stabilization compared to SV2A in its apo state, as well as to SV2A complexes bound with LEV-UCB1244283 or UCB1244283. Since labeled LEV is commercially available but BRV is not, we performed our binding experiments with ^3^H-LEV in the presence of UCB1244283. We assessed LEV binding using saturating concentrations of ^3^H-LEV, both with and without UCB1244283. We found that UCB1244283 induces a ∼2-fold increase in specific counts, consistent with previous functional experiments (Fig. 1c)^29^. Increased LEV binding upon addition of UCB1244283 enabled measurement of the half maximal effective concentration (EC_50_) of UCB1244283 binding, which is 0.6 ± 0.1 µM for wild-type SV2A (Fig. 1d). We also measured the binding affinity of LEV in the presence of saturating concentrations of UCB1244283, finding that LEV binds with a K_d_ of 3.8 ± 0.8 µM for wild-type SV2A in agreement with earlier studies^29^ (Fig. 1e).

### Cryo-EM of the SV2A-BRV-UCB1244283 complex

Our thermostability and previous binding data indicated that the SV2A-BRV-UCB1244283 complex exhibits the highest thermostability and affinity^30^, consequently we selected this complex for our structural studies. We purified SV2A together with LEV or BRV and UCB1244283 by previously established methods along with 8783-Nb to stabilize the SV2A lumenal domain (LD)^21^. Guided by our functional analysis, we used saturating concentrations of LEV or BRV and UCB1244283 throughout the SV2A purification. The purified SV2A complexes were subsequently analyzed by single particle cryo-EM, yielding a low-resolution map of the SV2A-LEV-UCB1244283 complex, and a 3.05 Å map of the SV2A-BRV-UCB1244283 complex after local refinement focusing on the TMD (Fig. 2a, S1). The refinement of the SV2A-BRV-UCB1244283 complex enabled the accurate modeling of all the residues in the TMD and intracellular domain (ICD) (Fig. S2). Notably, we observed two distinct density features in our map, one for BRV at the primary site and another for UCB1244283 at a site located ‘above’ the primary binding site towards the lumenal vestibule (Fig. 2a). When viewed from the lumenal side of SV2A, the binding of BRV and UCB1244283 appears to be positioned approximately in the middle of N-terminal and C-terminal TMDs, with a slight preference towards the C-terminal TMD, indicating that in the lumenal facing conformation these ligands bind SV2A predominantly with residues located in the C-terminal TMD. Furthermore, UCB1244283 binding to the allosteric site appears to directly block the dissociation of BRV from the primary site (Fig. 2b).

**Fig. 2.**
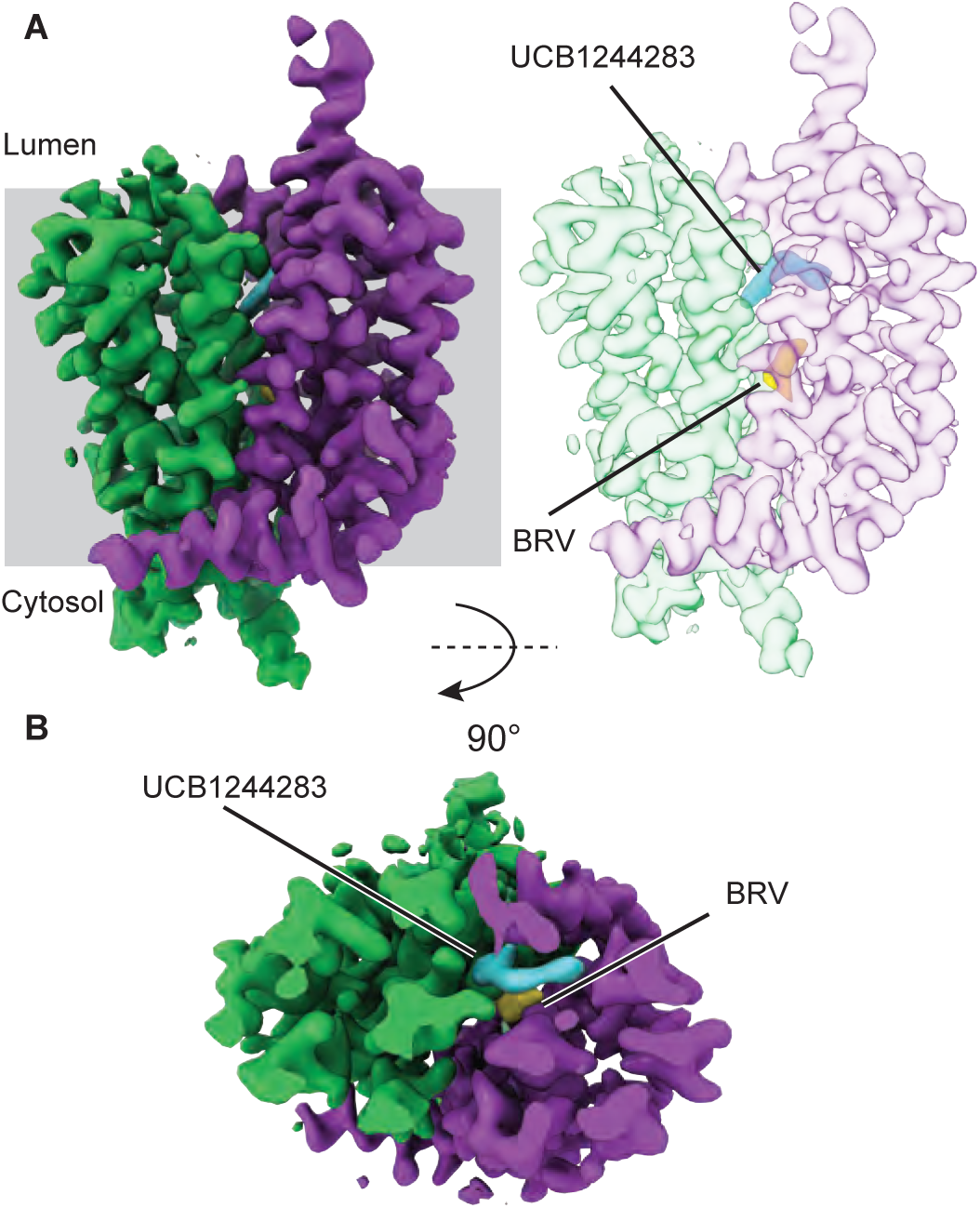
Structural overview of the SV2A-BRV-UCB1244283 complex. **A.** Cryo-EM reconstruction of the SV2A-BRV-UCB1244283 complex showing. The side view of the structure, showing transmembrane helices TM1-6 (green) and TM7-12 (violet). UCB1244283 (cyan) binds within a lumenal vestibule above BRV (yellow), oriented toward the SV lumen. **B.** View of the SV2A-BRV-UCB1244283 complex from the lumenal side, highlighting the binding site of BRV and UCB1244283. The density of SV2A has been sliced to allow viewing of the ligands. The binding site of both ligands is located between the two halves of SV2A.

### Molecular architecture of the UCB1244283 and BRV binding sites

BRV and UCB1244283 were modeled into their corresponding density features in our cryo-EM map. (Fig. 3a,b). Consistent with recent findings, the pyrrolidone group of BRV interacts with the primary binding site through hydrophobic interactions between Trp300 and Trp666, while the carbonyl of the pyrrolidone hydrogen bonds with Tyr462. The aromatic residues, Trp300, Trp666, and Tyr462 have been identified in several studies as key residues for binding both LEV and BRV^21,22,31,32^. The propyl group attached to the pyrrolidone extends into a subpocket near Tyr461 and TM10. Additionally, the amide group of BRV interacts with Asp670, playing a crucial role in the binding of both LEV and BRV, while the attached ethyl group forms van der Waals interactions with Leu176, Ile273, Phe277, and Cys297, which are important residues for racetam binding^21,22^. A group of conserved charged residues that are present in SV2 family members are found in the N-terminal TMD near the primary binding site: Arg262 on the lumenal end of TM4, Asp179 and Glu182 in TM1, and we observe that these residues coordinate density features for putative water molecules in our structure (Fig. S3).

**Fig. 3.**
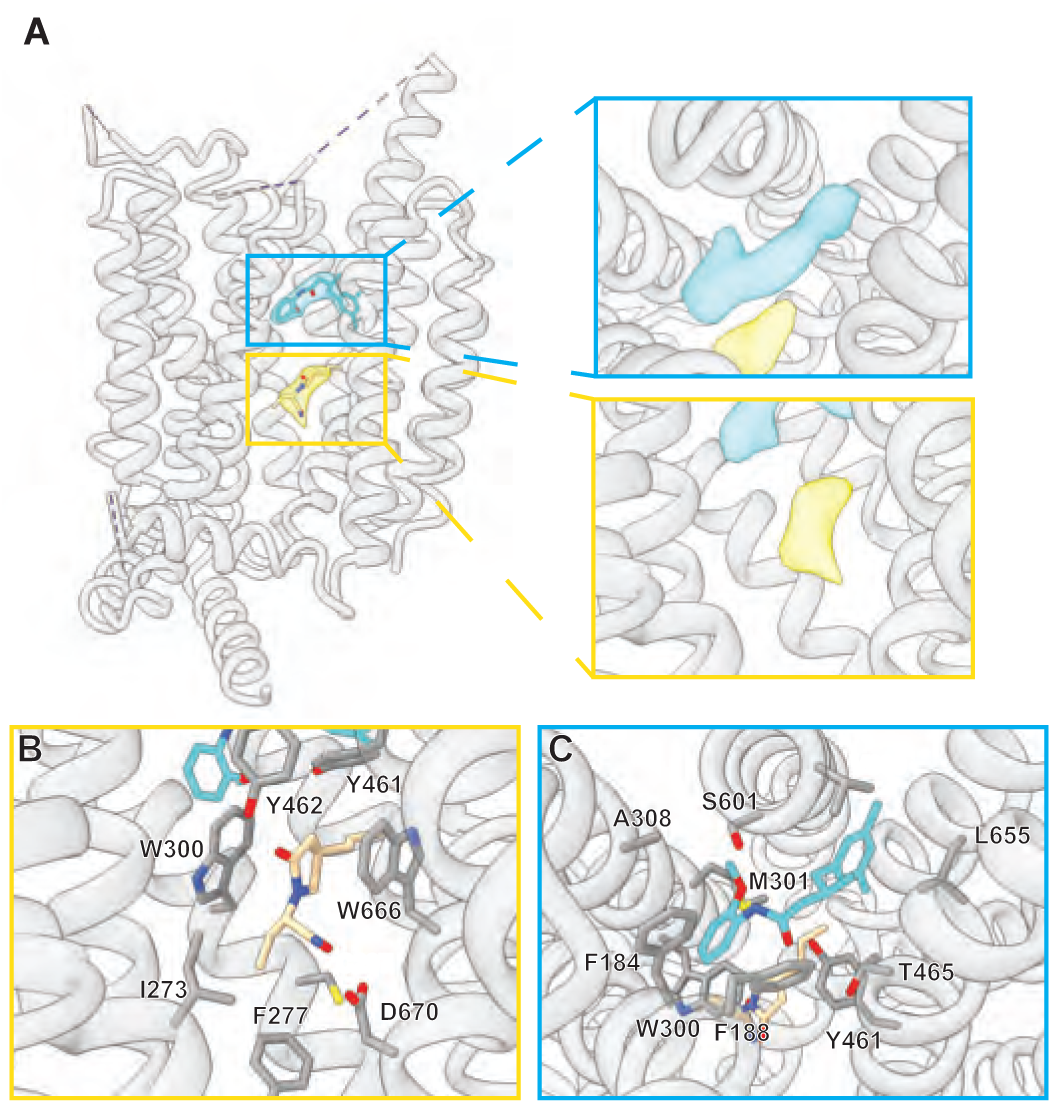
Architecture of the BRV and UCB1244283 binding sites. **A.** The side view illustrates the positioning of BRV and the R-enantiomeric form of UCB1244283 within the cryo-EM density, occupying the primary (yellow) and allosteric (cyan) binding sites, respectively. The inset (right) highlights the density features associated with BRV and UCB1244283. **B**. A close-up view of the BRV (tan sticks) binding site, with interacting residues shown as gray sticks. **C.** A close-up view of the UCB1244283 (cyan sticks) binding site, with key residues involved in binding shown as gray sticks.

The binding site of UCB1244283 is located in a lumenal vestibule between the two halves of the TMD (Fig. 3a,c), primarily constituted by residues from TM1, 5, 7, and 10. Since UCB1244283 is a racemic mixture of both *R*- and *S*-enantiomers, we modeled both isomers into the ligand density. The *R*- and *S*-forms fit equally well to the density (correlation coefficient: 0.79 vs. 0.77; Fig. S4), so it is possible that both may bind and therefore, we have included both fits in our PDB model but have primarily focused our description and analysis on *(R)*-UCB1244283 since the interactions of each isomer are similar. In TM1, Phe184 and Phe188, and Val597 in TM8 cap the upper part of the binding site on the lumenal side, positioned ‘above’ the 2-methoxyphenyl group of UCB1244283. Trp300 and Met301 in TM5 block access from the ‘lower’ part of the allosteric side to the primary binding site, while Ala308 is in the lumenal half of TM5, near Phe184 and Val597. Ser601 in TM8 is positioned near the oxygen of the methoxy group may also form a hydrogen bond with the amide (NH) moiety of UCB1244283 while Leu655 in TM10 is located adjacent to the dimethylphenyl group. TM10 contributes several key residues for UCB1244283 binding, with Thr465 and Tyr461 forming a polar network that interacts with the oxygen of the amide group in UCB1244283 (Fig. 3c).

### Analysis of the allosteric site amongst SV2 family members

To understand if UCB1244283 can bind other members of the SV2 family, we compared the sequence of the UCB1244283 binding site in SV2A with those of SV2B and SV2C (Fig. 4a). We also compared our structure with the SV2B-PSL complex and the AlphaFold-predicted^33^ structure of SV2C (Fig. 4b,c). Phe184 and Phe188 are absolutely conserved across the SV2 family members, while several other key residues involved in UCB1244283 binding are divergent. In brief, the residue equivalent to Ala308 in SV2A is a serine in both SV2B and SV2C, and Thr465 is conserved as a threonine in SV2B and replaced by a serine in SV2C. We observed more substantial differences near the dimethylphenyl group, where the equivalent residue to Leu655 in SV2A is a glutamine in SV2B and a leucine in SV2C. Finally, Gly659 which straddles the primary and allosteric sites and contributes to a subpocket in the primary site for larger ligands, such as UCB-2500, is a cysteine in SV2B and an asparagine in SV2C. Superposition of structures shows that the larger side chains of the asparagine and cysteine in SV2C and SV2B, respectively, would likely occupy the allosteric site near the dimethylphenyl group of UCB1244283.

**Fig. 4.**
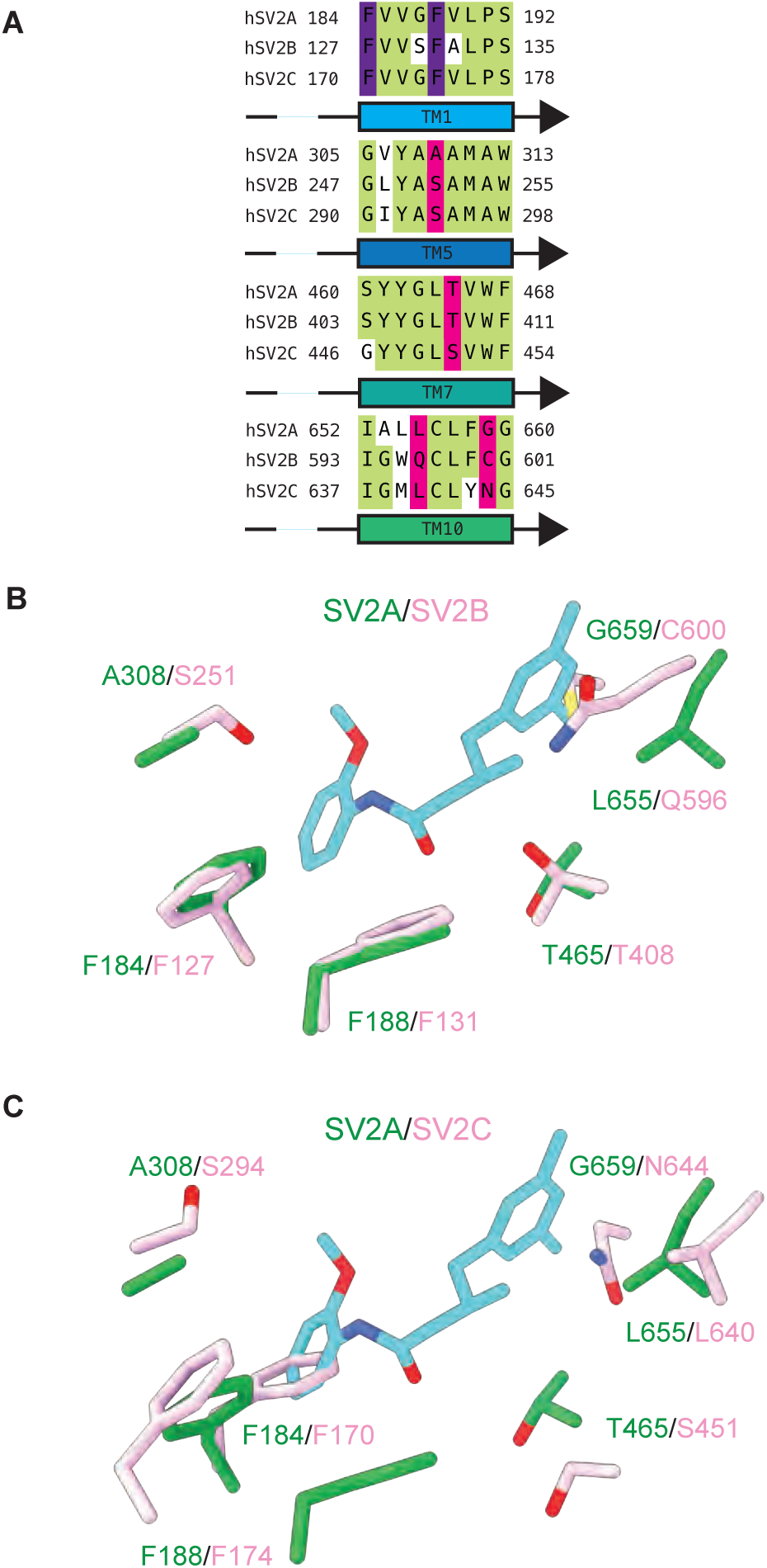
Structural comparison of the UCB1244283 binding site and selectivity for SV2A. **A.** Alignment of residues around the allosteric binding site of SV2A, SV2B and SV2C. Conserved residues are highlighted in green, residues directly involved in binding are shown in purple, and pink indicate residues directly involved in binding that are not absolutely conserved. **B.** Superposition of SV2A (green) and SV2B (pink) residues in the UCB1244283 binding site. **C.** Superposition of SV2A (green) and SV2C (pink) residues in the UCB1244283 binding site.

### Molecular basis of binding and selectivity of UCB1244283

To investigate the molecular determinants of UCB1244283 binding and selectivity, we studied the effects of mutants in the UCB1244283 binding site. We used two different strategies for mutating residues: first, residues which are predicted to be important for binding and are invariant or highly conserved amongst all SV2s were mutated to alanine, this includes F184A, F188A, and T465A; and second, residues which differ amongst SV2s were mutated to the corresponding residue in either SV2B or SV2C, which includes A308S, L655Q, G659C, and G659N. We then determined the binding affinity by measuring the EC_50_ for UCB124483 binding for each mutant (Fig. 5a).

**Fig. 5.**
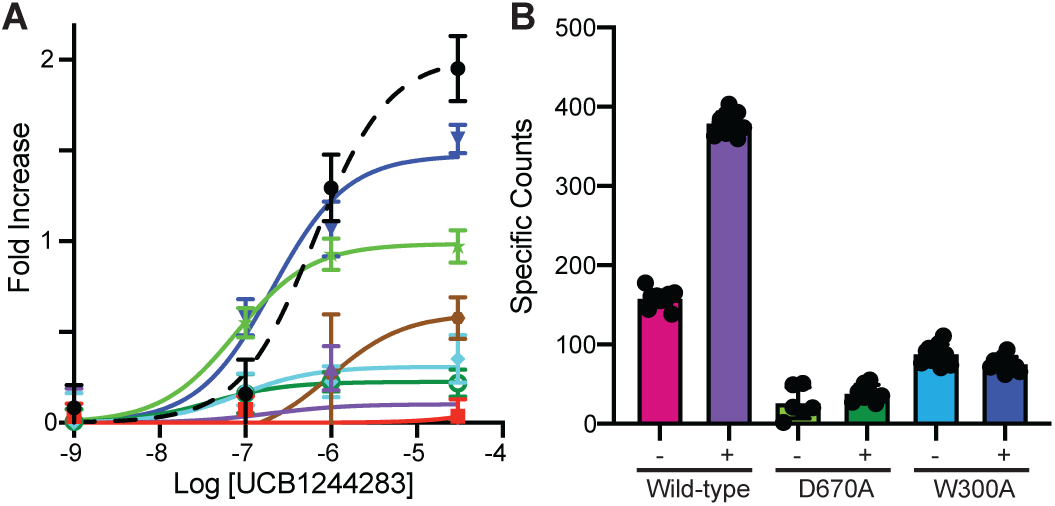
Molecular determinants of UCB1244283 binding and the effect of primary site mutations on the UCB1244283-induced enhancement of LEV binding. **A.** Concentration-response curves illustrating the effects of UCB1244283 on the binding of ^3^H-LEV to SV2A and allosteric-binding site mutants. UCB1244283 was used at concentrations of 30 µM, 1 µM, 100 nM and 1 nM, and incubated with ^3^H-LEV (3.3 µM) in the presence of SV2A and each mutant to determine the EC_50_ values in triplicate. The measured EC_50_ values were 0.6 ± 0.1 µM for wild-type SV2A (black circles) and 0.20 ± 0.04 µM, 0.07 ± 0.01 µM, and 1.0 ± 0.8 µM for SV2A-F184A (blue inverted triangles), SV2A-F188A (lime stars), and SV2A-G659C (brown hexagons) mutants, respectively. EC_50_ values for A308S (green open circles), T465A (cyan diamonds), L655Q (purple triangles), and G659N (red squares) could not be confidently determined by fitting due to their low activation binding counts. Data is shown as mean **±** SEM (n = 7). **B.** UCB1243283 (30 µM) binding does not enhance the maximum specific binding (B_max_) of ^3^H-LEV (3.3 µM) to SV2A-D670A and SV2A-W300A mutants.

For many of the mutants that we analyzed, a substantial reduction in the B_max_ was observed which precluded accurate EC_50_ fitting. The binding data demonstrates that A308S, L655Q, T465A, and G659N mutants do not show a significant change in LEV binding. The G659C mutant by contrast displayed a ∼4-fold reduction in B_max_ but exhibited a similar EC_50_ for UCB1244283 compared to wild-type SV2A. Finally, the F184A and F188A mutants showed minor reductions in B_max_ and exhibited enhanced EC_50_ values for UCB1244283 by ∼3- and 9-fold, respectively.

Previous studies suggest that UCB1244283 acts differently on the BRV and LEV complexes. UCB1244283 slows association and dissociation rates of BRV and increases its K_d,_ while LEV binding increases the B_max_ by 2-fold without significantly affecting LEV binding kinetics^28–30^. These observations led to the hypothesis that UCB1244283 binding may expose a second LEV binding site. To address this hypothesis, we measured LEV binding in the presence of UCB1244283 using the W300A and D670A mutants, which are known to disrupt LEV binding at the primary binding site^21,22^. In these experiments, we were unable to measure any significant LEV binding with either mutant with or without UCB1244283 (Fig. 5b).

### UCB1244283 and UCB-2500 stabilize a locked conformation of TMD

We compared our SV2A-BRV-UCB1244283 structure to SV2A bound to UCB-2500 and SV2B bound to PSL^21^, our superpositions revealed that the fluorines of the trifluorobutyl and chlorodifluoroethyl groups in UCB-2500 or PSL are located within ∼4 Å of the dimethylphenyl group in UCB1244283 (Fig. 6a,b).We compared the TMDs and ICDs between the SV2A-BRV-UCB1244283 complex and other SV2A structures^21,22^, bound with UCB2500, BRV, LEV, and the SV2B-PSL complex (Table 1), finding that the conformation of the SV2A-BRV-UCB1244283 complex most closely resembles the SV2A-UCB2500 complex. Comparison of the SV2A-BRV-UCB1244283 structure with the LEV-bound (PDB: 8JS8) or BRV-bound (PDB: 8K77) SV2A structures revealed substantial differences in the conformation of the luminal half of TM1 (Fig. 6c). In the BRV- and LEV-bound structures of SV2A the TM1 helix is substantially longer with a kink, featuring a 3_10_ helix spanning residues 181-184. In contrast, this region is a continuous α-helix and is shorter in the SV2A-UCB2500, SV2B-PSL, or SV2A-BRV-UCB1244283 structures (Fig. 6c,d, Fig. S5a-e). In the SV2A-BRV-UCB1244283 structure, this α-helical region undergoes a helical displacement of 4.7 Å (Cα to Cα distance for Gly187) compared to the SV2A-BRV structure. This conformational change results in blocking of both the allosteric and primary binding sites by Phe188. In the SV2A-BRV structure, Pro191 forms a hinge point after the 3_10_ helix and residues 191-197 are α-helical, whereas in our SV2A-BRV-UCB1244283, SV2A-UCB-2500, and SV2B-PSL structures, this region is modeled as a structured loop. Comparison of residues in the primary binding site show several other significant changes between various complexes, primarily in the positioning of Trp300 and Met301 which are shifted toward the primary binding site in the SV2A-BRV-UCB1244283 and SV2A-UCB-2500 complexes *vs.* the SV2A-BRV and SV2A-LEV complexes (Fig. 6e). We also observe shifts in the lumenal halves of TM2, 5, 7, 8, 10, and 11 in our SV2A-BRV-UCB1244283 and SV2A-UCB-2500 structures relative to the BRV-bound SV2A structure (Fig. 6f, Fig. S5f-i). These TMs contribute the majority of the residues responsible for binding both ASMs and UCB1244283, and these rearrangements modify the helical packing between TMs. They also alter the hydrogen bonding network in TM8, which is more extensive in the SV2A-BRV-UCB1244283 complex, involving Lys621, Asn667, Ser294, Asn612, Thr605, and Met301 (Fig. S5j-m). In TM8, we observe UCB-2500 or BRV-UCB1244283 binding to SV2A induces the formation of a 3_10_ helix, spanning residues 603-609, in both structures. Notably, binding of UCB1244283 and BRV to SV2A leads to repositioning of specific residues, including Ser601 and Thr605 in TM8 and Leu655 in TM10, compared to other SV2A structures, including SV2A-UCB2500 (Fig. 6g). These alterations facilitate the remodeling of allosteric site that allows UCB1244283 binding. Interestingly, the residues Thr465, Pro469, Val608, and Cys656, which are in proximity to the repositioned residues show no significant changes. (Fig. 6g,h).

**Fig. 6.**
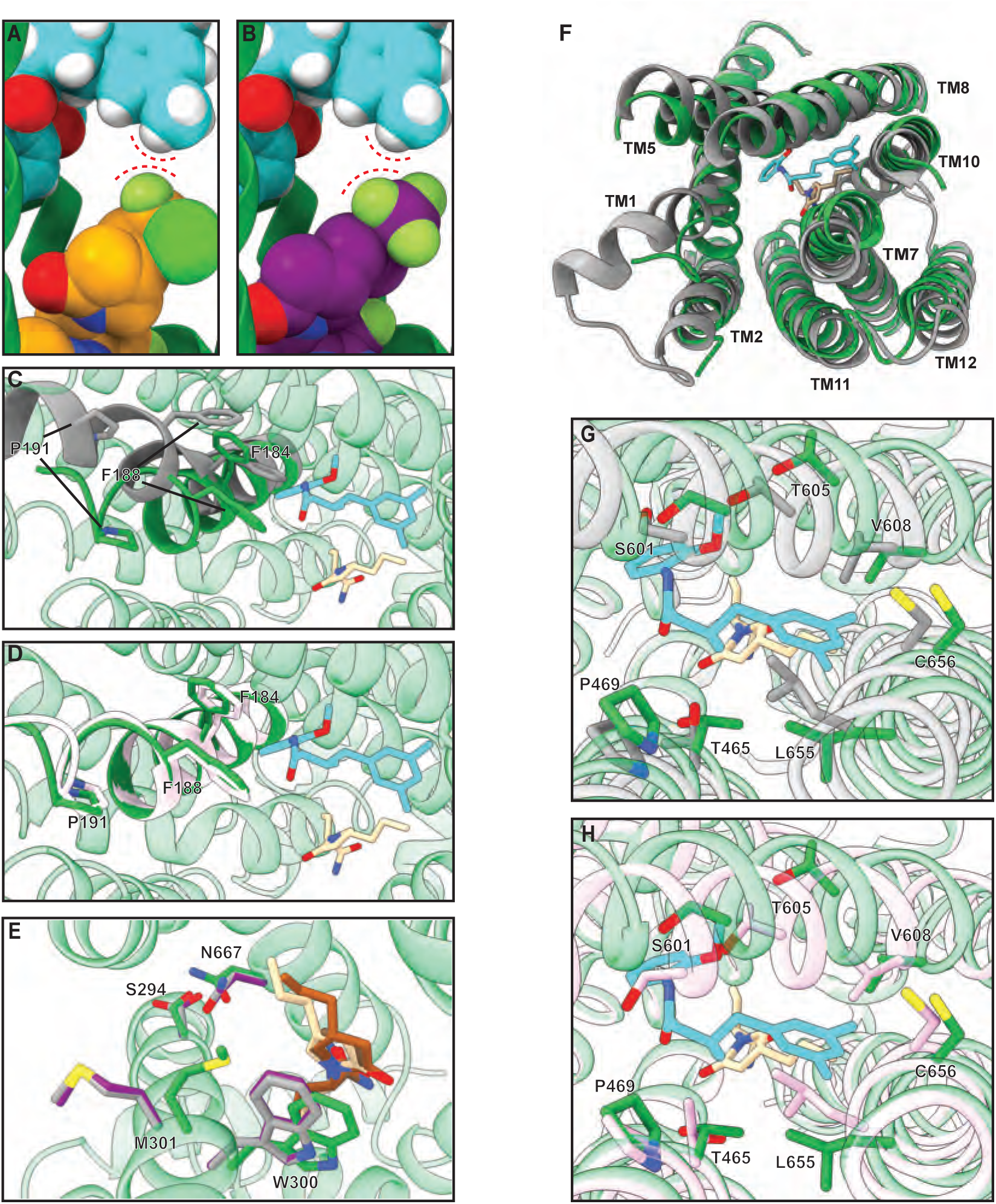
Conformation differences between the SV2A BRV-UCB1244283 and other ligand complexes. **A.** Sterics of the UCB1244283 (carbons, blue; hydrogens, white) and UCB2500 (orange) ligand binding sites, compared using the SV2A-BRV-UCB1244283 and SV2A-UCB2500 (PDB: 8UO9) complexes. Dotted lines indicate atoms that are predicted to be sterically incompatible. **B.** Sterics of the UCB1244283 and PSL (purple) binding sites compared using the SV2B-PSL complex (PDB: 8UO8). These models demonstrate that the presence of UCB2500 or PSL at the primary binding site sterically obstructs UCB1244283 binding to the allosteric site. **C.** Structural alignments of the SV2A-BRV (PDB: 8K77) and SV2A-BRV-UCB1244283 complexes reveal a conformational transition, from a 3_10_-helix to α-helix, in the TM1 helical region near the UCB1244283-binding site. **D.** Structural alignments of the SV2A-BRV-UCB1244283 and SV2A-UCB2500 (PDB: 8UO9) complexes show the same conformation of TM1. **E.** The structural superposition between the SV2A-BRV-UCB1244283 complex (green sticks, BRV shown in tan) and the SV2A-BRV (grey sticks, BRV shown in light brown) or SV2A-LEV (purple sticks, LEV shown in dark brown) structures revealed alterations in primary site residue positioning. **F.** Comparison of the SV2A-BRV (PDB: 8K77) and SV2A-BRV-UCB1244283 complexes highlight conformational rearrangements in the transmembrane helices surrounding the primary and allosteric binding site observed from the lumenal side. **G.** Comparison of the allosteric site between SV2A-BRV-UCB1244283 (green) and the SV2A-BRV structures (grey). The position of BRV and UCB1244283 are shown in tan and blue sticks respectively. **H.** Comparison of the allosteric site between SV2A-BRV-UCB1244283 (green) and the SV2A-UCB-2500 structures (pink).

**Table 1.** Comparison of RMSD values between SV2A complexes. The TMD and ICD of each complex were compared to the SV2A-BRV-UCB1244283 complex.

| Complex name | RMSD ($\text{\AA}$) |
| --- | --- |
| SV2A-UCB2500 (PDB: 8UO9) | 1.4 |
| SV2A-LEV (PDB: 8JS8) | 2.0 |
| SV2A-BRV (PDB: 8K77) | 2.0 |
| SV2B-PSL (PDB: 8UO8) | 2.8 |

## Discussion

The SV2A-BRV-UCB1244283 complex traps SV2A in a conformation where both the primary and allosteric binding sites are occupied by ligands, allowing for detailed mechanistic insights into allosteric modulation of ASM binding. UCB1244283 binds to a lumenal site in a vestibule located in the pathway which leads to the primary binding site, where ASMs like LEV and BRV bind (Fig. 7a). The allosteric site is positioned ‘above’ the primary binding site, directly obstructing the dissociation of ASMs from the primary binding site (Fig. 7b). Therefore, our structure provides a direct explanation of the mechanism by which UCB1244283 slows the dissociation rate of ASMs and increases their binding affinities. The molecular basis for how MFS transporters are allosterically controlled by small-molecule binding is also not well understood. While allosteric sites have been identified in other MFS transporters, they typically bind either lipid, detergent, or ions^34^. The allosteric site in SV2A is a novel binding site for small-molecule ligands and our results suggest that similar mechanisms and binding sites may be present in other SV2 family members and related MFS transporters. This allosteric mechanism closely parallels that of the unrelated serotonin transporter (SERT), where allosteric ligands similarly influence the binding of inhibitors to the primary site^35^.

**Fig. 7.**
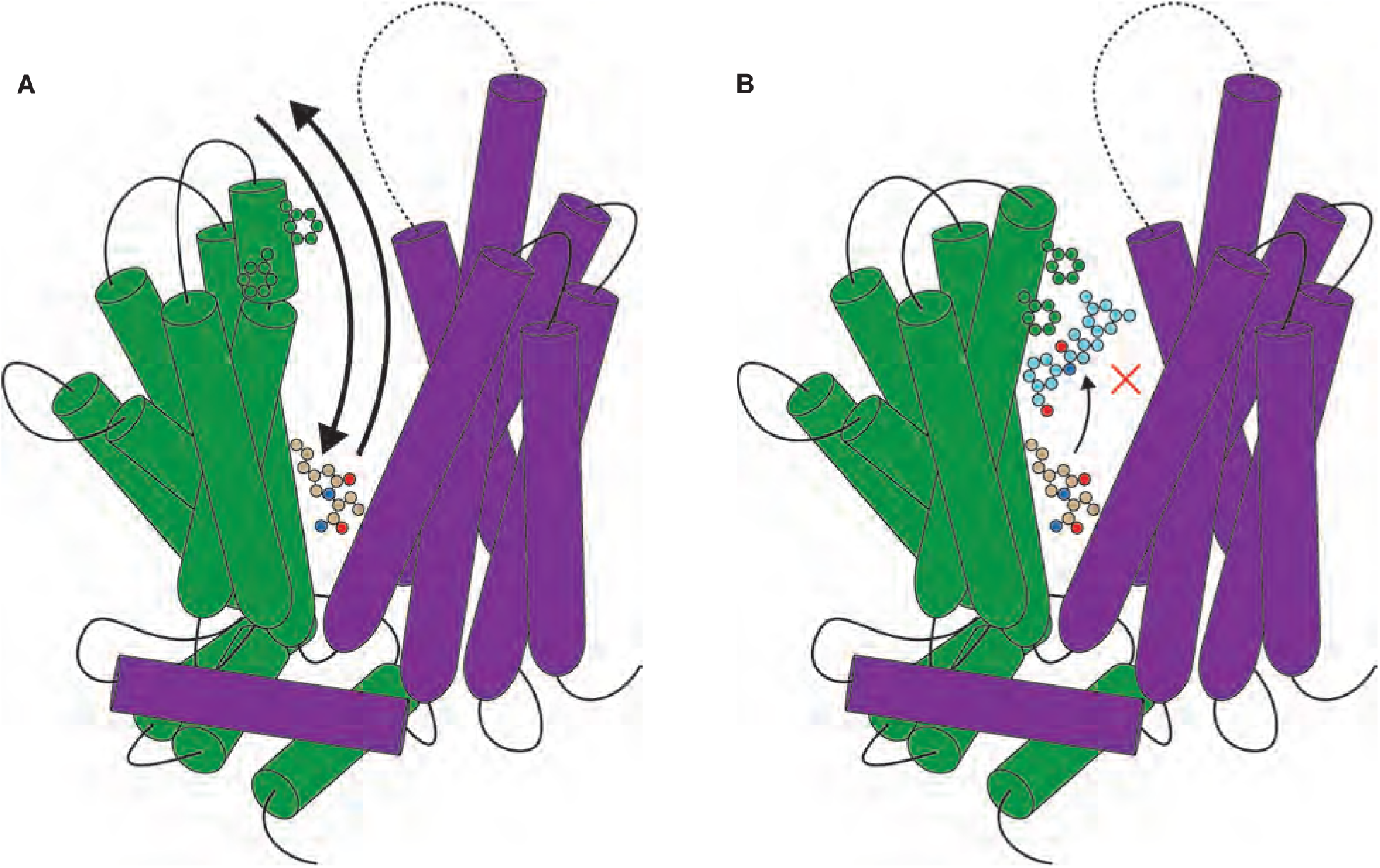
Model illustrating the effects of the SV2A allosteric modulator UCB1244283 on the enhanced binding kinetics of antiseizure medications. **A.** SV2A is composed of two six-transmembrane (TM) bundles, N-terminal (TM1–6, green) and C-terminal domains (TM7–12, purple). Structural analyses of SV2A-BRV (PDB: 8K77) and SV2A-BRV-UCB1244283 reveal a single binding site for BRV, located between the two halves of the transporter. The TM1 helix, towards the lumenal side, is oriented away from the C-terminal domain due to a kink created by the 3_10_-helix, which results in an open upper lumenal gate. **B.** UCB1244283 binding to the allosteric site stabilizes BRV at the primary binding site by sterically obstructing its unbinding, while inducing conformational changes in the surrounding TMs, notably, inducing a conformation change in the lumenal half of TM1 from a 3_10_-helix to an α-helix in TM1, resulting in reorientation of TM1 toward the C-terminal half of the TMD. Key phenylalanine residues lining the allosteric site are shown in green. This conformation results in closure of the upper lumenal gate in the SV2A-BRV-UCB1244283 structure, thereby enhancing the binding affinity of BRV at the primary binding site.

The molecular features of the UCB1244283 binding site are predominantly composed of hydrophobic and uncharged polar residues, which allow for the binding of molecules like UCB1244283. By contrast, the primary binding site is formed by hydrophobic and charged side-chain residues, which accommodate the binding of more polar ASMs, such as LEV, BRV and PSL^21–24^ (Fig. 3b,c). UCB1244283, when administered in combination with UCB30889, a LEV analogue, exhibits protective effects against both tonic and clonic convulsions, but these effects are observed exclusively following intracerebroventricular administration^28^. This property is thought to be due to the poor blood-brain barrier passage of UCB1244283, which limits its effectiveness as an oral medication^21–24^. These insights into both the primary and allosteric binding sites could aid in developing molecules with enhanced specificity and superior pharmacokinetics, resulting in more effective targeting of SV2s. Given the close proximity of these sites, several strategies have been employed in other transporters, GPCRs, and other proteins to effectively target neighboring binding sites, which may be applicable to SV2s^36–40^. For example, in SERT, bivalent serotonin molecules connected by PEG spacers have been developed to bridge both sites^37–39^. These ligands are potent inhibitors but may not have significant potential for therapeutic development due to their size and other pharmacological properties. Another potential approach is the development of high-affinity versions of molecules like UCB1244283 that target the allosteric site of SV2A with optimized pharmacological properties that could be used in conjunction with existing therapies to improve potency and specificity. Finally, bitopic ligands have been designed in GPCRs to target both allosteric and orthosteric sites; these ligands are functionally selective which may allow for improved efficacy^38–40^. It is conceivable that such strategies may also prove useful in further development of novel molecules against SV2s and other related SLC transporters.

The cryo-EM density observed at the allosteric site allowed fitting of both (*R)*- and (*S)*-enantiomers of UCB1244283, indicating that at the current resolution, it is not possible to resolve whether there is preferential binding of one isomer over the other (Fig. S4). Therefore, further investigations are required to clarify the enantiomeric specificity of the allosteric site in SV2A. EC_50_ measurements with (*R)*- or (*S)*-UCB1244283 would provide direct insight into whether one enantiomer binds with higher affinity or if there is no preference.

Previous saturation binding studies have indicated that UCB1244283 differentially affects BRV binding to SV2A compared to LEV^28–30^. In brief, UCB1244283 increases BRV binding primarily by increasing its affinity and B_max_, while it increases LEV binding mainly through an increase in B_max_. This observation has led to the hypothesis that the number of LEV binding sites increases from one to two, suggesting that LEV and BRV either bind to different sites or modulator induces significant conformational changes in SV2A, altering its interactions with BRV or LEV. Mutagenesis of residues around the primary binding site (W300F, F277A, G303A, F658A, Y462A, W666A, I663A, D670A, and V661A) reduced the binding of both BRV and LEV^22,30–32^. However, for the I273A, K694, and S294A mutants, the modulator’s effect on increasing the B_max_ for LEV was abolished, but not for BRV, further suggesting that SV2A ligands interact distinctly with the primary or other binding sites^28–30^. Alignment of the primary binding site in the SV2A-BRV-UCB1244283 complex with the structures of SV2A-BRV (PDB 8K77), and SV2A-LEV (PDB 8JS8), shows what appear at first glance to be only minor changes, primarily around Trp300, Asn667, and Met301, while Ile273, Lys694, Ser294 superimpose closely across all complexes (Fig. 6e). Our binding experiments with W300A and D670A mutants, together with UCB1244283, suggest that there is not a second LEV binding site (Fig. 5b). Rather, we hypothesize that subtle differences in the conformation of primary binding site residues and/or the surrounding TMs are responsible for the previously observed binding differences between the UCB1244283-BRV and -LEV complexes (Fig. 6c,e,f). Consistent with this hypothesis, the sidechain positions of Trp300 and Asn667 are shifted, with Met301 oriented toward the primary site in the SV2A-UCB1244283-BRV complex, in contrast to facing the lipid membrane as seen in the SV2A-LEV and SV2A-BRV complexes^22^. Trp300 and Met301 reside in the same region of the binding site as Ile273 and Lys694, sitting ‘beneath’ the allosteric site. The methoxyphenyl ring of UCB1244283 pi-stacks with the sidechain of Trp300, and this interaction may be partially responsible for repositioning of this residue in comparison to other SV2A-drug complexes. We propose that these residues are part of an allosteric network that connects the primary site to the allosteric site and is responsible for transmitting the binding differences that are observed between the UCB1244283-LEV and BRV complexes^28–30^ as well as the SV2A-LEV and SV2A-BRV complexes without UCB1244283^22^.

Superposition of the SV2A-BRV-UCB1244283 structure with other SV2 structures such as the SV2A-BRV, -LEV, -UCB-2500 or SV2B-PSL complexes, demonstrates that there are no large-scale conformational changes which occur upon occupancy of the allosteric site by UCB1244283. However, our structural analysis highlights several critical conformational changes in the primary binding site, allosteric site, and the lumenal half of the TMD, that distinguish the SV2A-BRV-UCB1244283 complex from other previously characterized complexes. Comparison of the SV2A-BRV-UCB1244283 and SV2A-UCB-2500 structures with SV2A-BRV and SV2A-LEV revealed significant conformational differences in the upper luminal part of TMDs, particularly in TM1 and TM8, which result in a locked TMD conformation. In addition to the blockade of the primary site by UCB1244283 binding, the conformational changes in TM1 involving Phe184 and Phe188 further obstruct the lumenal side of the allosteric site in both the SV2A-BRV-UCB1244283 and SV2A-UCB-2500 complexes, which likely also prevents primary ligand dissociation. The enhanced EC_50_ values for F184A and F188A mutants are likely a result of further stabilization of this locked TMD conformation, which is driven by TM1. We hypothesize that the replacement of bulky phenylalanine sidechains with alanines either stabilizes the α-helical conformation of TM1 or promotes UCB1244283 association by enhancing local flexibility and reducing steric hinderance. This may facilitate structural adjustments that allow more favorable interactions for UCB1244283, such as with the oxygen of Thr465. We speculate that residues involved in an N-terminal TMD polar network may play a role in proton binding and coupling to transport. The putative water densities observed in this region are unique to our SV2A-BRV-UCB1244283 structure (Fig. S3). Changes in the position of primary site residues in the SV2A-BRV-UCB1244283 and SV2A-UCB-2500 complexes also alter the hydrogen bonding network in TM8 relative to the SV2A-BRV and SV2A-LEV complexes, and these changes appear to be transmitted from the primary site to the lumenal half of the TMD, thereby repositioning key residues in the allosteric site. Notably, in the BRV-UCB1244283 bound SV2A structure, we observed that TM8 forms a hydrogen bond with Ser294, a feature not observed in other SV2A structures (Fig. S5). Previous binding data showed that the S294A mutation reduced the affinity of BRV and LEV and abolished the effect of UCB1244283 for LEV but not for BRV, demonstrating that BRV and LEV bind to closely related sites but interact in distinct ways^28^. In TM8, we observed that Asn612 forms a hydrogen bond with Cys297 in both SV2A-BRV and SV2A-LEV structures^22^. We have previously shown that Cys297 is involved in selective binding of LEV, by mutating the corresponding glycine in SV2B to a cysteine (G240C), the inhibition constant (K_i_) for LEV is increased by 100-fold for SV2B G240C, with no significant effect on BRV^21^. In our SV2A-BRV-UCB1244283 structure, Asn612 hydrogen bonds to Ser294, instead of Cys297 when UCB1244283 binds, suggesting that this interaction network may also be involved in selective binding of ASMs (Fig. S5j-l). Therefore, we propose that these networks control the conformation of the TMD, and these conformational changes can be regulated by either primary or allosteric ligand binding, thus explaining why the SV2A-BRV-UCB1244283 and SV2A-UCB-2500 complexes are more closely related to one another *vs*. SV2A-BRV and SV2A-LEV complexes.

We also focused on several non-conserved residues in the allosteric site, analyzing mutants at these positions using EC_50_ measurements to gain insights into the selectivity of UCB1244283 for SV2B and SV2C. In SV2B, the equivalent residues to Leu655 and Gly659 in SV2A are glutamine and cysteine, respectively. We predict that the L655Q mutant in SV2A would induce a steric clash with the dimethylphenyl group, while the G659C mutant may still accommodate UCB1244283 in the allosteric site. Although the G659C mutant exhibited a similar EC_50_, the B_max_ was significantly lower, and since the L655Q mutant showed no significant EC_50_ activation with UCB1244283, we conclude that UCB1244283 does not bind SV2B. For SV2C, the most notable difference in the allosteric site is the replacement of the equivalent residue to Gly659 with asparagine. The longer side chain of the asparagine, compared to glycine or cysteine in SV2A or SV2B, suggests that the G659N mutant would also disrupt UCB1244283 binding. Our binding data confirms that the G659N mutant also displayed minimal modulation of LEV binding with UCB1244283. Future studies will be required with wild-type SV2B and SV2C to understand whether UCB1244283 is an SV2A-specific allosteric modulator. UCB1244283 is known to selectively bind to the allosteric site in the presence of certain primary site ligand complexes, while binding to others, such as, PSL and likely UCB2500, are not compatible^15^. Our structural analysis provides an explanation for this selective binding behavior. We find that the chloro-difluoro and trifluoro groups of these ligands would likely create steric hinderance by interfering with the dimethylphenyl group of UCB1244283, thereby disrupting its binding (Fig. 6a,b). In addition to LEV and BRV, UCB1244283 also allosterically modulates the binding kinetics of UCB30889, a higher-affinity analogue of LEV containing a larger azidophenyl group^28^. At present, it is not known whether UCB1244283 can bind SV2A in combination with other important ligand complexes, such as UCB-J^41^ or SDI-118^42^ and this will require further investigation with ^3^H-labeled ligands.

UCB1244283 may serve as a useful tool for trapping endogenous molecules at the primary site, aiding the investigating of SV2A function^43^. SV2s have been thought to be transport proteins but the endogenous substrate has not yet been identified^9,10,14,44–46^. For most transporters, the substrate affinity lies typically in the micromolar range, which makes purifying substrate-bound transporters, especially from native sources, challenging^34^. Transported substrates are predicted to bind to the primary site^34^, and since UCB1244283 blocks unbinding from this site, we predict that it could be used to facilitate the isolation of SV2A bound to its natural substrate in this conformation^21,22,26,43,44^. We also speculate that endogenous ligands could bind to the primary and the allosteric sites of SV2A to modulate it’s activity ^34^.

In conclusion, this study provides a detailed structural analysis of the essential synaptic vesicle protein, SV2A, bound to the ASM, BRV and an allosteric modulator, UCB1244283. We demonstrate how UCB1244283 binds in a hydrophobic pocket, only in combination with LEV and BRV, and selectively binds to SV2A within the SV2 family. BRV and LEV bind to a shared primary site but interact distinctly with SV2A. Binding experiments with mutants at the primary binding site in the presence of UCB1244283 suggest that SV2A does not contain a second cryptic LEV binding site, which was previously proposed to become accessible upon UCB1244283 binding^28–30^. We further highlight the role of conserved residues in the human SV2 family, which are not directly part of the primary or allosteric sites but contribute to a network involved in mediating crosstalk between these two functional sites. These differences were primarily localized to the luminal regions of the TMD, particularly in TM1 and TM8, which resulted in a locked TMD conformation in our structure. The binding of UCB1244283 to the allosteric site induces changes in primary site residues which alters the local conformation of the TMD. These structural changes are transmitted between both sites, affecting the conformation of key residues and facilitating ligand interactions. Notably, the enhanced binding affinity of BRV in the SV2A-BRV-UCB1244283 complex is likely driven by these changes and by obstruction of the primary site by UCB1244283. Collectively these interactions are distinct from those observed in the SV2A-LEV or SV2A-BRV complexes^22^ without UCB1244283 and explain the mechanism of positive allosteric modulation of ASM binding.

## Materials and Methods

### Small-molecule ligands

PSL, ^3^H-LEV, and UCB1244283 were obtained from UCB Pharma through a Material Transfer Agreement. ^3^H-LEV was purchased through this agreement from American Radiolabeled Chemicals (ART 1832). BRV (Cat No. R020192) and LEV (Cat No. L8668) were purchased from MuseChem and Sigma, respectively.

### Construct design and cloning

The SV2A construct used in this study has been described previously and behaves similar to full-length SV2A in ligand binding experiments^21^. In brief, the first 64 amino acids were deleted from the 5’ end of human *SV2A*, and the 3’ end was extended with sequences coding a 3C protease site (L-E-V-L-F-Q-G-P), mVenus^21^, TwinStrep, and 10-His-tags. This sequence was cloned into pEG-BacMam^47^, resulting in *SV2AΔ64-mVenus* construct. The *SV2AΔ64-mVenus* construct was subject to site-directed mutagenesis to create F184A, F188A, A308S, T465A, L655Q, G659C, and G659N mutants. The mutants were verified by sequencing.

### Fluorescence-detection size-exclusion chromatography

Fluorescence-detection size-exclusion chromatography (FSEC) analysis^48,49^ was conducted using a Shimadzu HPLC instrument equipped with a refrigerated autosampler, multi-wavelength detector, and a Superose 6 Increase 5×150 column. The fluorescence of mVenus (excitation at 515 nm and emission at 528 nm) was monitored in whole cell lysates solubilized using 10 mM lauryl maltose neopentyl glycol (LMNG) detergent and 1 mM cholesteryl hemisuccinate (CHS) for mVenus-tagged SV2A. For purified proteins, tryptophan fluorescence (excitation at 280 nm and emission at 325 nm) was monitored.

### Expression and purification of 8783-Nb

The expression and purification of 8783-Nb has been described previously^21^. In brief, 8783-Nb expression was carried out in the periplasm of BL21-DE3 *E.coli* cells from pET26-8783-Nb plasmid harboring a PelB sequence. The periplasmic fraction of the cells was prepared by incubating the harvested cells with ice-cold hypertonic solution (20mM Tris-HCl, pH 8.0, 20% sucrose, 1 mM EDTA)^50^ for 30 minutes, followed by centrifugation at 7600 g. The 8783-Nb was purified from the diluted supernatant by binding to a Ni^2+^-NTA gravity flow column, followed by size-exclusion chromatography (SEC) of eluted fractions using a Superdex 75 Increase 10/300 column (Cytiva).

### SV2A Expression and purification of SV2A

The recombinant expression of the SV2AΔ64-mVenus protein was carried out in mammalian tsA201 cells using the baculovirus-mammalian (BacMam) cell expression system, using Sf9 cells to produce baculovirus^47^. Harvested cell pellets were resuspended in TBS_150_ (20 mM Tris-HCl pH 8.0, 150 mM NaCl), protease inhibitor cocktail (200 nM aprotinin, 2 µM leupeptin, 2 µg/mL pepstatin) and were lysed using high-frequency ultrasonic waves (Misonix Sonicator 3000). Unbroken cells and cell debris were removed by centrifugation at 10,000 rpm (SS34 rotor, Sorvall RC6) for 10 minutes, followed by ultracentrifugation of supernatant fraction at 40,000 rpm (Ti45 rotor, Beckman Coulter) for 90 minutes at 4°C to isolate membranes. Cell membranes were resuspended in TBS_150_ and incubated with 30 µM UCB1244283 and 100 µM of levetiracetam (LEV) (Sigma, Cat No. L8668) or brivaracetam (BRV) (MuseChem, Cat. Number R020192) for 30 minutes on ice, followed by solubilization using 10 mM lauryl maltose neopentyl glycol (LMNG) and 1 mM cholesteryl hemisuccinate (CHS) at 4°C for 90 minutes. The insolubilized fraction was removed by ultracentrifugation at 40,000 rpm for 60 minutes at 4°C, and the cleared supernatant was loaded onto a gravity-flow column containing 4 ml GFP-nanobody conjugated sepharose resin. The resin was washed with 20 column volumes of TBS_150_ containing 0.02 mM LMNG, 0.020 mM CHS, 0.050 mM glyco-diosgenin (GDN), 30 µM UCB1244283 and 10 µM of LEV or BRV. The resin bound with SV2AΔ64 was incubated overnight with 3C protease to remove mVenus, TwinStrep, and 10-His-tags. The next day, the flow-through containing SV2AΔ64 were mixed with 4-5 molar excess of 8783-Nb, followed by concentration to 500 µl using a 100 kDa concentrator (Amicon). The SV2AΔ64-8784nb complex was injected onto a Superose 6 10/300 column (Cytiva) in TBS containing 0.02 mM LMNG, 0.020 mM CHS, 0.050 mM GDN, and 30 µM UCB1244283 and 25 µM LEV or 10 µM of BRV. SEC fractions of the homogenous peak corresponding to monomeric SV2AΔ64 were further individually analyzed by FSEC.

SEC fractions that were homogenous based on FSEC analysis were pooled and concentrated to 5.3 mg/ml or 8 mg/ml using a 100 kDa concentrator before addition of 30 µM UCB1244283 and 1 mM of LEV or BRV, respectively. The concentrated sample was ultracentrifuged at 60,000 g prior to cryo-EM grid preparation.

### Cryo-EM sample preparation and data collection

2.5 µl of SV2A was applied to glow discharged Quantifoil holey carbon grids (1.2/1.3 200 mesh copper). Grids were blotted for 2-4 seconds with a blot force of 2-4 (100% humidity, 4°C) using an FEI Vitrobot MK IV (ThermoFisher). SV2A grids were imaged at the University of Pittsburgh on a Titan Krios G3i operating at 300 kV equipped with a Falcon 4i direct electron detector and a Selectris energy filter set to a slit width of 10 eV. Movies were collected at a pixel size of 0.719 Å/pixel with defocus ranges from −0.5 to −2.0 µm with a total dose of 50 e/Å^2^.

### Cryo-EM data processing and model building

A total of 7,888 exposures were selected from 10,356 micrographs, and CryoSPARC v4.2^51^ was used to perform patch motion correction and contrast transfer function (CTF) estimation. Particles were picked using reference-free blob picker (∼1.9 million particles) followed by particle extraction at a box size of 336 pixels. These particles were binned to 128 pixels and classified through five rounds of hetero-refinement, using ‘decoy’ classes with random density, empty detergent micelles and low-resolution ab-initio models of SV2A, followed by 2D classification. The selected particles were used to generate an ab-initio model to create projection-based templates. Next, we performed a patch motion correction on a random subset of 100 exposures, which were then used to generate a denoise model. This model was subsequently utilized in denoising the entire set of micrographs, which were used for particle picking (∼ 3.6 million particles) using the template picker. Particles from blob pick and template pick were individually classified multiple times (10-15x) using hetero refinement as described above followed by several rounds of 2D classification. Duplicate particles were removed that resulted in 185,565 particles. These particles were used for generating three 3D-maps through ab-initio reconstruction. One 3D-reconstruction consisting of 78,706 particles was chosen for re-extraction at a box size of 256 pixels, which were refined to 3.44 Å after non-uniform refinement followed by local refinement using the TMD mask. These particles were re-extraction at a box size of 336 pixels and were subjected to two rounds of Bayesian polishing in RELION v.3.1^52,53^. Polished particles were subjected to two rounds of 3D-classification in cryoSPARC that separated a set of 36,^21^7 particles, which were refined to 3.05 Å after local refinement using the TMD mask. SV2AΔ64 (PDB: 8UO9) model was fit into the SV2A-BRV-PAM cryo-EM density maps, which was manually modelled in Coot^54,55^, iteratively real space refined in Phenix (version 1.20.1)^56^, and validated by comparing the half maps and refined model.

### Thermostability assays

SV2AΔ64-mVenus containing tsA201 cell membrane were solubilized in TBS_150_ containing 5 mM LMNG, 1 mM CHS, with and without ligands (apo, 20 µM BRV, 30 µM UCB1244283, and BRV/UCB1244283 at the same concentrations) for 60 minutes at 4°C. These samples were ultracentrifuged at 60,000 g for 30 minutes to remove cell debris and the supernatant was heated at various temperatures for 15 minutes, followed by ultracentrifugation for 20 minutes to remove aggregates. The supernatants were subjected to FSEC^47^ on a Superose 6 Increase 5×150 column for detection of mVenus fluorescence.

### Radioligand binding assays

SV2AΔ64-mVenus wild-type or single residue mutant plasmids were transfected into tsA201 cells in suspension culture (200 ml) using polyethylenimine 40K (PEIMAX). After 48-60 hours post transfection, cells were harvested, and membrane enriched with wild-type or each mutant were prepared as described above. Protein expression of each construct was analyzed by FSEC, and membranes containing equal amounts of proteins were used in the filter binding assays. We prepared a working radioligand ^3^H-LEV by diluting concentrated ^3^H-LEV (200 µM, 5 Ci/mmol) using unlabeled LEV to a final concentration of 2.5 µM and 100 µM, respectively.

For saturation-binding studies, SV2AΔ64-mVenus enriched membranes were incubated for 2 hours in presence of 30 µM UCB1244283 and a 2-fold serially diluted ^3^H-LEV (10 µM to 0.0781 µM) in triplicate. Specific binding of ^3^H-LEV to SV2AΔ64-mVenus or mutant membranes was calculated by subtracting nonspecific binding from the total binding. Nonspecific binding was measured by residual binding observed in presence of 10 mM LEV.

For EC_50_ calculations, SV2AΔ64-mVenus or single residue mutant enriched membranes were incubated for 45 minutes on ice with ^3^H-LEV at 3.3 µM and a series of UCB1244283 ligand concentrations at a final concentration of 30 µM, 1 µM, 100 nM and 1 nM, respectively. Specific binding was calculated by subtracting nonspecific binding from the total binding, which was measured by residual binding observed in the presence of unlabeled 1 mM of padsevonil (PSL). For all experiments, incubated samples were filtered on glass microfiber filters (Cytiva, Cat No. 1822-6580, 25mm) presoaked in 1% polyethylenimine (PEI), followed by a very rapid washing with 20 ml ice-cold TBS_150._ Filters were dried and were transferred into a 24-well plate (PerkinElmer, Cat No. 1450-402) followed by the addition of 500 µl of scintillant to each well for measuring radioligand binding. Counts were measured using a Microbeta-2 scintillation counter (PerkinElmer). The K_d_ and EC_50_ values were calculated by GraphPad Prism 10 software using non-linear fitting for one site-specific binding and log (agonist) vs response (three parameters) equation, respectively.

## References

ACEHOLDER FOR INSERTION OF FINAL REFERENCE LIST.

## Acknowledgments

We thank James Conway and the Pittsburgh Center for Cryo-EM for their support.

## Author contributions

Conceptualization: A.M., M.F.M., P.S.H., and J.A.C. Funding acquisition: J.A.C. Methodology: A.M., M.F.M., and J.A.C. Developed compounds and contributed chemical expertise: A.H., L.P. Contributed to the selection of compounds for SV2A and to map and pose interpretation: M.G., L.P., A.H., A.M., J.A.C., P.S.H. Investigation: A.M. Discussion of experiments: M.G. Confirmed the pose of the compounds: M.L., P.S.H. Characterized the compounds in vivo/in vitro: M.G. and C.W. Visualization: M.F.M., A.M., and J.A.C. Supervision: P.S.H. and J.A.C. Writing—original draft: A.M., M.F.M. and J.A.C. Writing—review and editing: A.M., M.F.M., C.W., M.G., A.H., M.L., L.P., P.S.H., and J.A.C.

## Data and materials availability

All data needed to evaluate the conclusions in the paper are present in the paper and/or the Supplementary Materials. EMDB and PDB under the accession codes 49758 and 9NTC, respectively were released in online repositories.

