## Supplementary for "Mechanisms Underlying Allosteric Modulation of Antiseizure Medication Binding to Synaptic Vesicle Protein 2A (SV2A)"

### Supplementary Materials

Anshumali Mittal *et al.*

**This PDF file includes:**

Figs. S1 to S5
Tables S1

### Supplementary Text

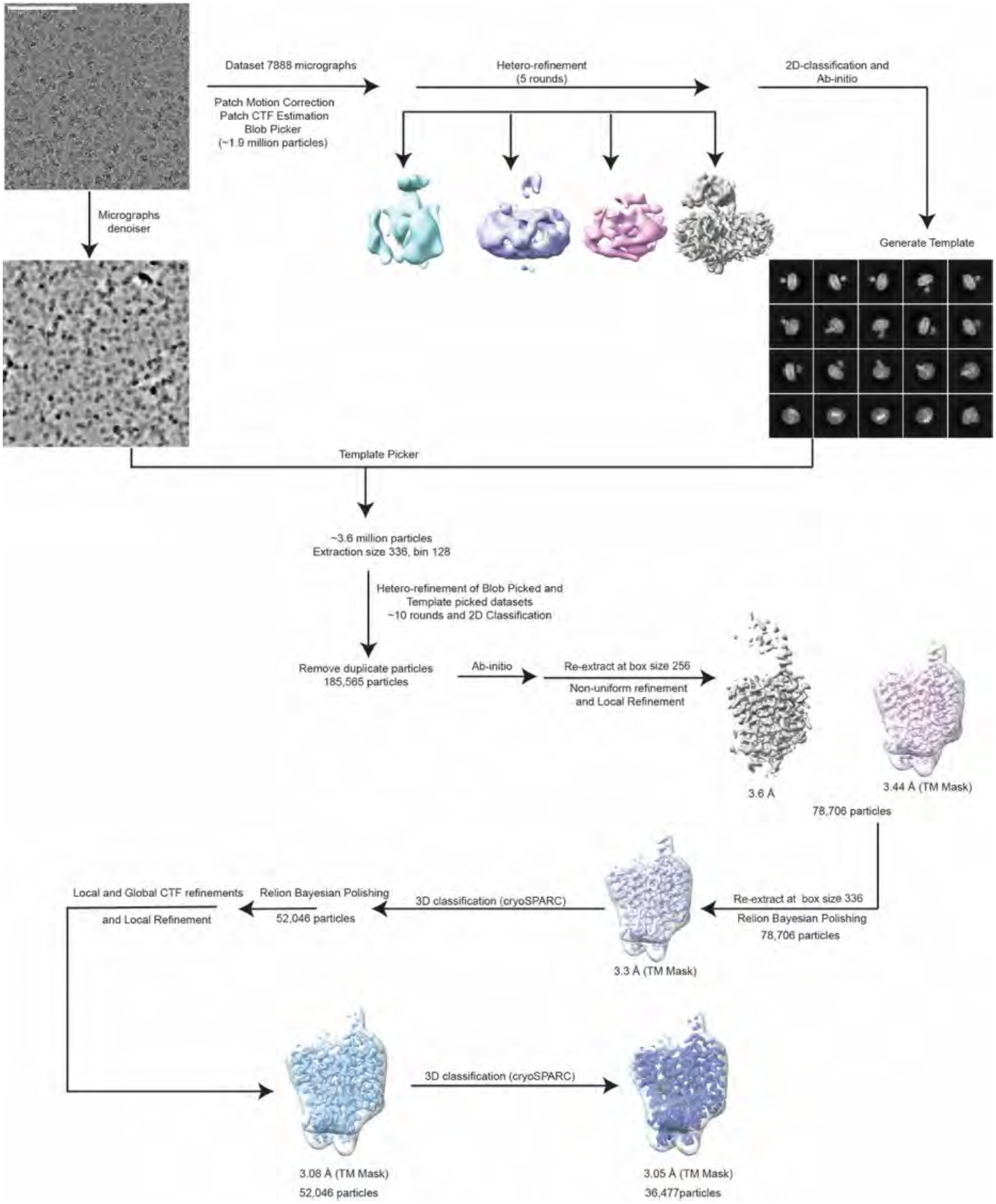

**Fig. S1.**

**SV2A-BRV-UCB1244283 cryo-EM data and processing**. A representative micrograph is shown with a scale bar of 100 nm. A total of 10,356 micrographs were collected and after patch motion correction and CTF estimation, blob picker was used to pick particles from manually selected 7888 micrographs (~1.9 million particles). These particles were initially classified by heterogenous refinement, using several ‘decoy’ volumes and one volume with SV2A features, followed by 2D classification. An initial volume was generated by Ab-initio reconstruction that was used for particle picking from 7888 micrographs using Template Picker (~ 3.6 million particles). Particles from Blob pick and template pick were individually classified using several rounds of heterogenous refinement and 2D Classification as mentioned previously. Duplicate particles were removed, and the remaining particle stack was analyzed by ab-initio reconstruction, non-uniform, and local refinement with a mask focused on transmembrane region. The resulting particle stack, 78706 particles, were reextracted at a full box size (336 pixels) and processed two times in Relion using Bayesian Polishing including a 3D-classifications in cryoSPARC. A final map was generated using 36,477 particles at a resolution of 3.05 Å.

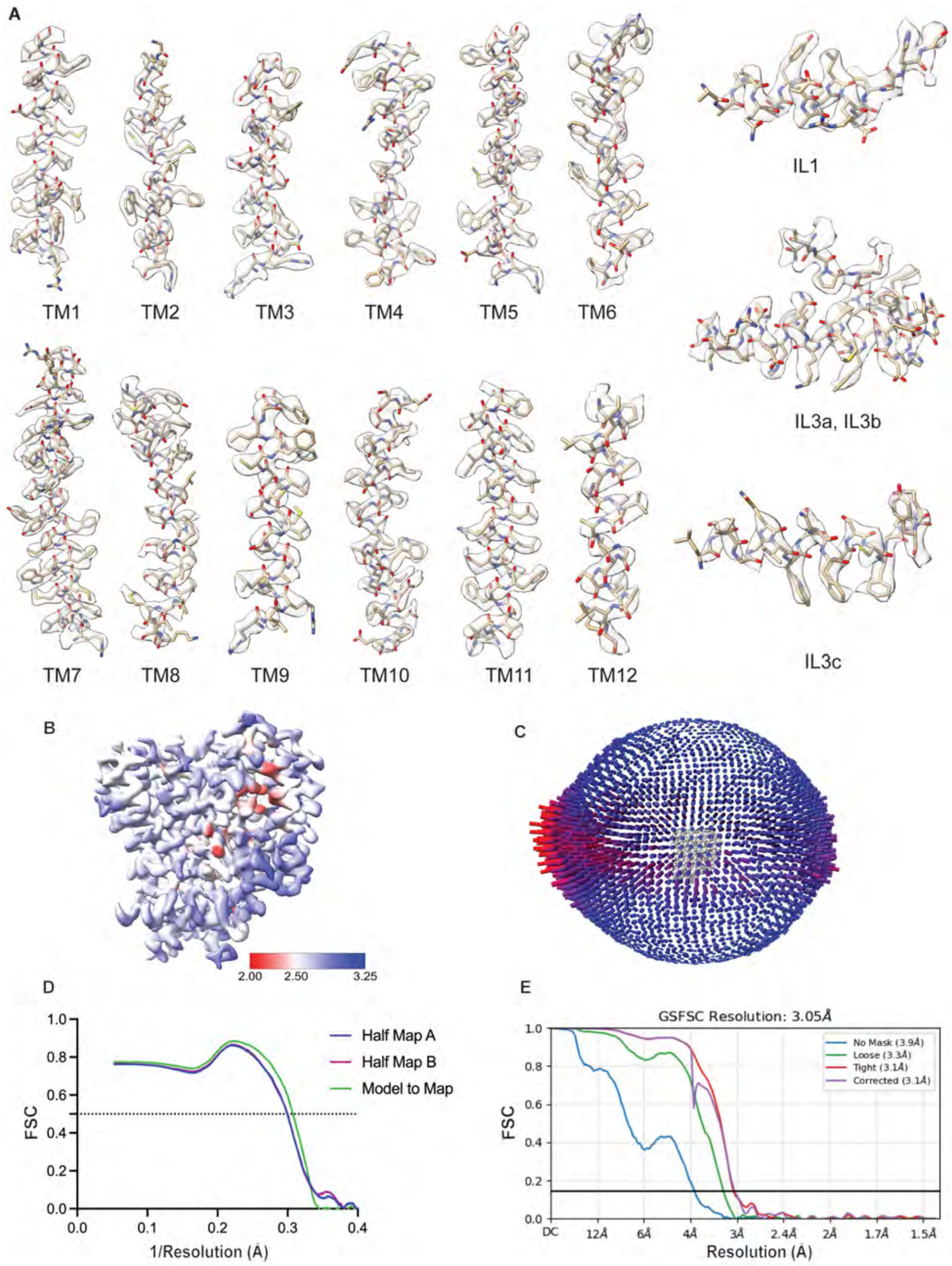
A
TM1
TM2
TM3
TM4
TM5
TM6
IL1
IL3a, IL3b
TM7
TM8
TM9
TM10
TM11
TM12
IL3c
B
2.00
2.50
3.25
C
D
FSC
1/Resolution (Å)
Half Map A
Half Map B
Model to Map
E
GSFSC Resolution: 3.05Å
No Mask (3.9Å)
Loose (3.3Å)
Tight (3.1Å)
Corrected (3.1Å)
FSC
Resolution (Å)
DC
12Å
6Å
4Å
3Å
2.4Å
2Å
1.7Å
1.5Å

**Fig. S2.**

**Interpretation of SV2A-BRV-UCB1244283**. **A.** Modelling of all 12 transmembrane helices, and the intracellular domain helices of SV2A. **B.** Local resolution of the SV2A transmembrane region map. **C.** Euler angle distribution of the SV2A transmembrane reconstruction. **D.** Map to model correlation of the SV2A model with the transmembrane SV2A. **E.** FSC curve of the SV2A transmembrane region map.

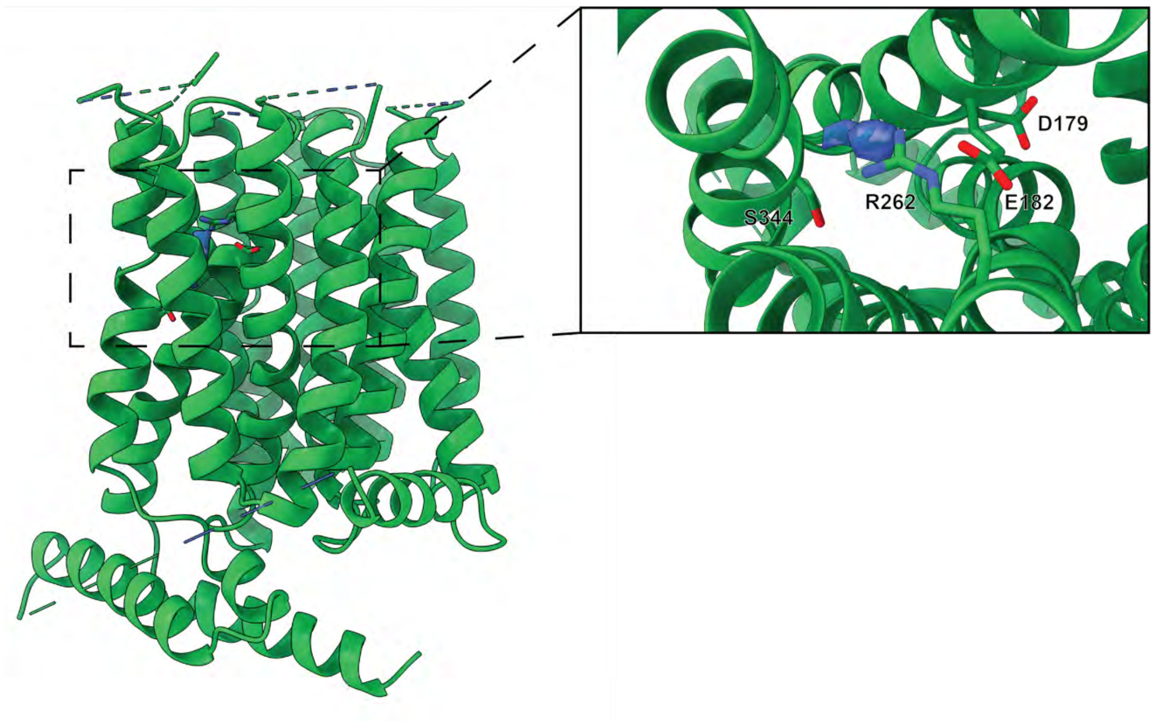

D179
S344
R262
E182

**Fig. S3.**

**Water network in the N-terminal TMD.** Densities for a putative water network involving Asp179 Glu182, Arg262, Ser344 were observed.

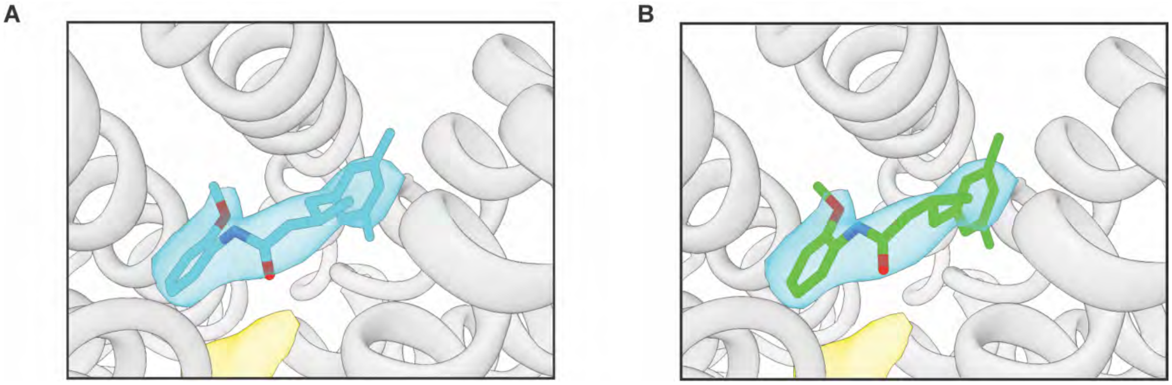
A
B

**Fig. S4.**

**Comparison of two poses of UCB1244283 in the cryo-EM density map**. **A.** Fit of the R-enantiomer. **B.** Fit of the S-enantiomer.

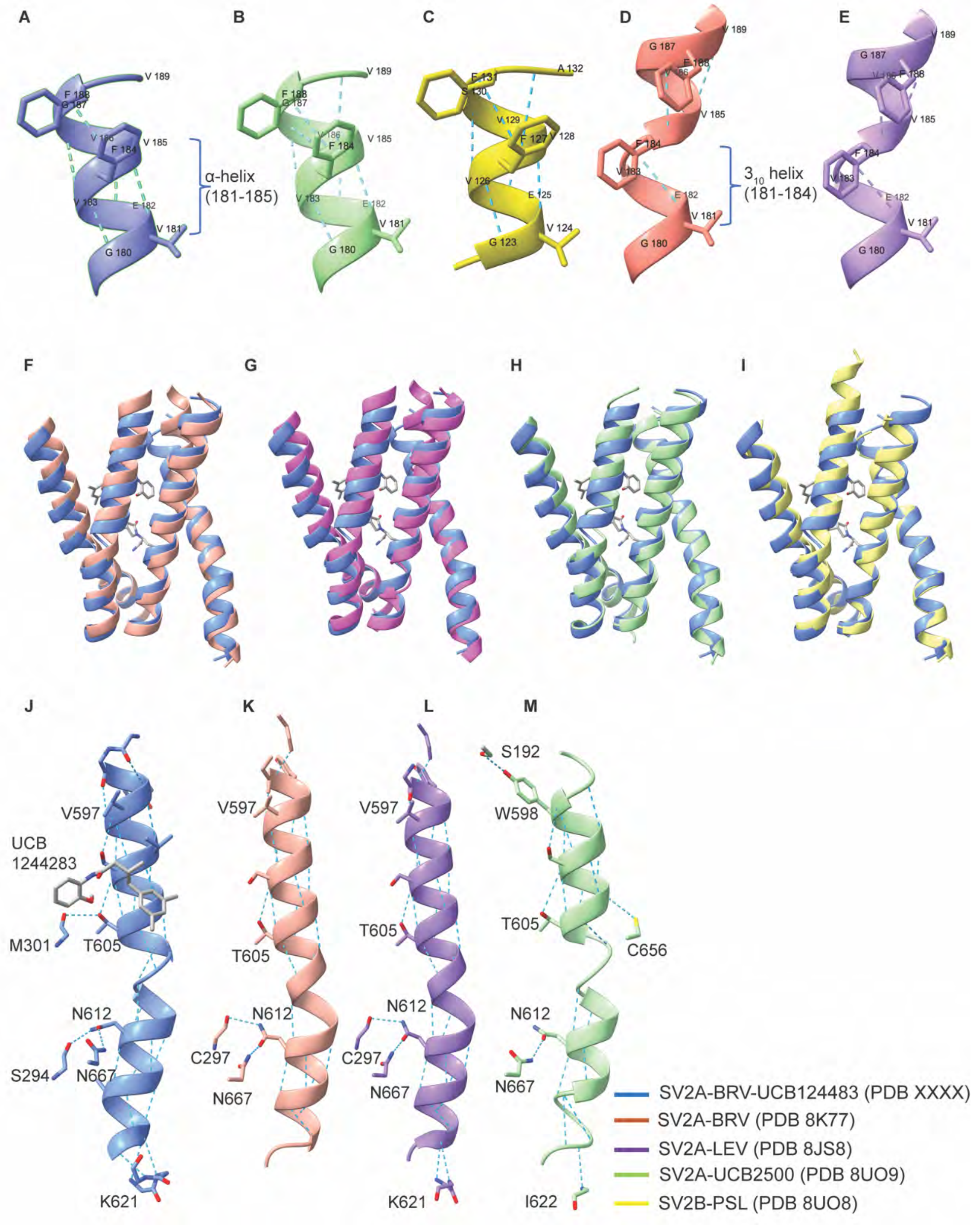
A
B
C
D
E
V 189
F 188
G 187
V 186
F 184
V 185
V 183
E 182
V 181
G 180
α-helix
(181-185)
A 132
F 131
S 130
V 129
F 127
V 128
V 126
E 125
V 124
G 123
3$_{10}$ helix
(181-184)
F
G
H
I
J
K
L
M
S192
V597
W598
UCB
1244283
M301
T605
C656
N612
S294
N667
C297
K621
I622
SV2A-BRV-UCB124483 (PDB XXXX)
SV2A-BRV (PDB 8K77)
SV2A-LEV (PDB 8JS8)
SV2A-UCB2500 (PDB 8UO9)
SV2B-PSL (PDB 8UO8)

**Fig. S5.**

**Comparison of conformational changes in TM1 and TM8 and their surrounding regions.** Comparison of secondary structure elements within TM1, residues 180-189. **A.** SV2A-BRV-UCB1244283 in blue. **B.** SV2A-UCB2500 (PDB 8UO9) in light green. **C.** SV2B-PSL (PDB 8UO8) in yellow. **D.** SV2A-BRV (PDB 8K77) in light coral. **E.** SV2A-LEV (PDB 8JS8) in violet. **F-I.** Superposition of TM1, TM5, TM8, and TM10 between SV2A-BRV- UCB1244283 in blue and SV2A-BRV in light coral or SV2A-LEV in violet or SV2A-UCB2500 in light green or SV2B-PSL in violet. **J-M.** Hydrogen bonding network of TM8, residues 594-619, with neighboring TMs.

**Table S1.**

Cryo-EM data collection, refinement and validation statistics.

| | SV2A-BRV- UCB1244283 (EMDB-XXXX) (PDB XXX) |
|---|---|
| **Data collection and processing** | |
| Magnification | 194,000x |
| Voltage (kV) | 300 |
| Electron exposure (e–/Å$^2$) | 50 |
| Defocus range (μm) | -0.5 to -2.0 |
| Pixel size (Å) | 0.719 |
| Symmetry imposed | C1 |
| Initial particle images (no.) | 5.5 million |
| Final particle images (no.) | 36,477 |
| Map resolution (Å) | 3.05 |
| FSC threshold | 0.143 |
| Map resolution range* (Å) | 2.5 – 3.25 |
| **Refinement** | |
| Initial model used (PDB code) | 8UO9 |
| Model resolution (Å) | 3.3 |
| FSC threshold | 0.5 |
| Model resolution range (Å) | 18.9 - 3.3 |
| Map sharpening | -86.8 |
| Model composition | |
| Non-hydrogen atoms | 6734 |
| Protein residues | 432 |
| Ligands | 4 |
| *B* factors (Å$^2$) | |
| Protein | 44.95 |
| Ligand | 45.72 |
| R.m.s. deviations | |
| Bond lengths (Å) | 0.04 |
| Bond angles (°) | 1.65 |
| Validation | |
| MolProbity score | 1.59 |
| Clashscore | 7.36 |
| Poor rotamers (%) | 0.3 |
| Ramachandran plot | |
| Favored (%) | 96.92 |
| Allowed (%) | 3.08 |
| Disallowed (%) | 0 |
